# Plasmonic imaging of living pancreatic beta-cell networks

**DOI:** 10.1101/2025.04.22.649945

**Authors:** Sidahmed Abayzeed, Daniel Galvis, Karen Regules Medel, Oscar Barajas Gonzalez, Kerry Setchfield, Michael G. Somekh, Kyle C. A. Wedgwood, Paul Smith

**Affiliations:** Faculty of Engineering, University of Nottingham; Centre for Systems Modelling & Quantitative Biomedicine (SMQB), School of Medical Sciences, University of Birmingham. Birmingham; Living Systems Institute, University of Exeter; EPSRC Hub for Quantitative Modelling in Healthcare, University of Exeter; School of Life Sciences, University of Nottingham

## Abstract

We present a novel plasmonic imaging technique for real-time, label-free tracking of bioelectrical interactions in live pancreatic beta-cell networks. Surface plasmon resonance microscopy (SPRM) is utilized to reveal synchronized glucose-induced intensity oscillations that are suppressed by calcium channel blockers. These oscillations are observed at the subcellular scale with a resolution of 1 μm. The technique can also uncover the extracellular spread of these oscillations beyond the cells. We further combine SPRM with network analysis to quantify coordinated electrical activity within the living cell network using both amplitude and phase-based metrics. Our results demonstrate a new method for studying electrical communication in pancreatic beta-cells, which could be crucial for understanding dysregulation in diabetes and advancing treatment development. This technique holds promise for investigating electrical connectivity in biological cell networks with applications in neuroscience, cardiac science, and bioelectricity in cancer, microbiology, development and regeneration.

**Graphical abstract:** 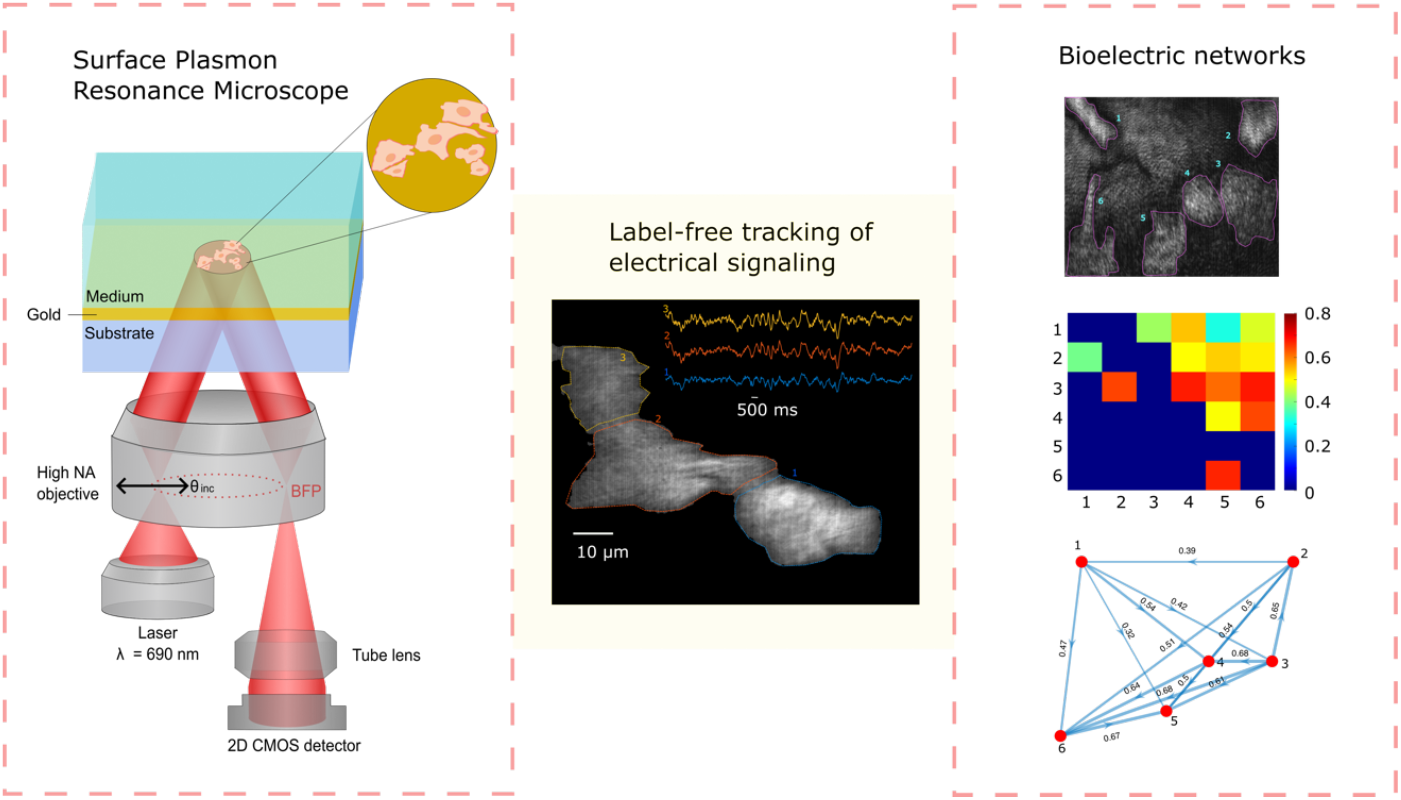

## Introduction

Biological cells have evolved complex molecular mechanisms for coordinated electrical signaling that are critical for a wide range of physiological and developmental processes. For example, neurons form networks to process sensory information ^1^ and coordinate the organism’s interactions with internal ^2^ and external ^3^ environments. Similarly, cardiomyocytes are electrically coupled via gap junctions to produce synchronous mechanical action ^4^. Hormone secreting cells also function in synchronized networks driven by coupled electric oscillations to produce secretions that are efficient for their function. For instance, pancreatic beta-cells, within the islet of Langerhans, are functionally connected ^5^, a feature crucial for regulated insulin release and glucose homeostasis ^5^. Furthermore, intercellular electrical communications via tunnelling nanotubules are reported in immune cells ^6^. More recently, novel oscillations of membrane potentials have been observed in human breast cancer cells with notable temporal correlations ^7^ - indicative of electrical coupling. Furthermore, a recent breakthrough revealed the role of electrical excitability in the progression of small-cell lung cancer ^8^. Similar to eukaryotic cells, prokaryotes coordinate metabolism via propagation of electrical signals in microbial biofilms ^9^. These examples highlight the significance of tracking electrical signals in live cells.

Electrical coupling in living biological cells is currently investigated via microelectrode arrays (MEAs) ^10^ and probes ^11^, which provide a direct measure of endogenous electrical signaling. However, MEAs and high density MEAs offer a limited spatial resolution (i.e. several micrometers) and low spatial sampling due the constraints of fabrication, electrical wiring and electrode spacing ^12^. Calcium and voltage imaging approaches are also employed to monitor synchronization of oscillations in intracellular calcium and transmembrane potential. However, these imaging approaches suffer from the drawbacks associated with fluorescence, such as limited temporal resolution and sensitivity. Although these technologies have driven a great advancement in our understanding of living cell networks ^13 14^, there is a need for new technology with enhanced capabilities that offer label-free, high-resolution and high-density characterization of electrical connectivity in living cells.

The development of label-free methods has allowed measurement of electrical signals *in vitro* and *in vivo*. For instance, plasmonic approaches have been applied to track action potentials in neurons ^15,16^ and cardiomyocytes ^17^. Furthermore, nitrogen vacancy diamond sensors ^18,19^, and graphene sensors ^20^ have also been introduced for label-free detection of electrical signaling in living neurons. We report a significant advancement in label-free microscopy: a technique that captures synchronized bioelectrical activity across living cell networks at a spatial resolution of 1 μm^2^. Unlike previous methods, our technique uniquely reveals how activity propagates through the extracellular space, extending analysis beyond direct cell-sensor contact. This capability opens new opportunities to investigate the role of the extracellular environment in bioelectrical signaling, with potential applications in wound healing, development, and regeneration. Furthermore, we merge surface plasmon resonance microscopy (SPRM) with network analysis techniques to explore the coordinated network activity and assess the temporal variations in connectivity.

SPRM was introduced in a pioneering work by Rothenhäusler and Knoll ^21^ and independently by Yeatman and Ash ^22^ in the late 1980s as a contrast enhancement technique achieved by illuminating biological samples, adhered to a gold or silver thin film, through the excitation of surface plasmons (SPs). Since then, SPRM has been investigated for studying cell adhesion, migration and proliferation in a label-free manner ^23^. Furthermore, several studies have demonstrated the ability of SPRM to reveal time-resolved processes ^24,25^ such as membrane biomolecular interactions ^26^ and exocytosis ^27^. SPRM of various cell types ^28^ such as neurons, cardiac and bacterial cells ^29,30^ have driven the interest of several groups globally; further demonstrating its diverse applications^31,32^.

The decades of research from several teams have innovated a variety of SPRM configurations and concepts, for instance, leveraging the change in phase ^33^ or intensity of reflected light around the surface plasmon resonance (SPR) position ^34^. This includes highly sensitive interferometric approaches that exploit the sharp phase transitions at the SP excitation angle ^35^. The high intensity gradients of the SPR curve are utilized for realizing widefield SPRM configurations ^36^. This is achieved by illuminating the sample at an angle of incidence with a non-zero and, ideally, a maximum gradient of SPR curve at a fixed or multiple azimuthal directions ^37^. The fixed-angle widefield configuration has been used to investigate cell-sensor interfaces ^38^ and realize impedance microspectroscopy of cells and biomolecules ^39,40^. The output of SPRM, such as changes in reflected light intensity at a high intensity gradient, probes the electric charge dynamics at the metal-electrolyte interface ^40^. This capability directly measures the electrical activity of living biological cells ^16^ due to the ionic perturbation of the double-layer capacitor at the cell sensor interface ^41^. We leverage this SPRM capability for fine-grained imaging of electrical activity from the sub-cellular level to a whole cellular network, which is anticipated to provide novel insights about the function of biological cell networks, with cutting-edge applications in cancer, diabetes, neuroscience and microbial colonies.

In this study, we used the pancreatic beta-cell line MIN6. MIN6 are a well-established, representative model, of primary beta-cells ^42^ and have been used in over 16,000 publications prior to 2025. They share the hallmarks of native beta-cells: glucose sensitive calcium-dependent action potentials, which are responsive to specific hormones and ion-channel drugs and are coupled to insulin secretion ^43^. Pancreatic beta-cells produce action potential electrical behaviour in response to an elevation in blood glucose ^44^. Briefly, a rise in plasma glucose inhibits the activity of ATP-sensitive potassium channels (KATP), which is predominantly responsible for their resting membrane potential, Vm of −70 mV. This leads to depolarisation of Vm and activation of voltage-gated Ca2+ channels (VGCCs) and calcium-dependent action potential (AP) electrical activity (+20 mV peak). The frequency of APs is graded with glucose concentration and degree of KATP block^44^, from this an emergent electrical behaviour of bursts or cluster of APs arises. Bursting consists of APs solely associated with depolarized, plateau Vm of ~ −40mV separated by periods of electrically silent hyperpolarized Vm of ~ −50mV ^45^. The molecular identity, expression profile, pharmacology, roles and regulation of KATP and the VGCCs within the pancreatic beta-cell are well documented ^46^.

## Results

### Surface plasmon resonance microscopy (SPRM) of pancreatic beta-cells

we present SPRM applied to the imaging of live cell networks, introducing the capability to monitor cellular interactions as well as the signal spread in their vicinity, for the first time. This is crucial to elucidate the propagation of information between cells embedded in a conductive extracellular environment. To achieve this, we fabricated SPR sensors based on gold thin films, as depicted in Fig. 1a, to excite SPs at the metal-dielectric interface. The resonance phenomenon is marked by a drop of reflectivity at a particular angle of incidence, termed SPR angle. Research has shown that this resonance position is sensitive to refractive index variations at the interface and our group has investigated SPR sensitivity to externally applied voltage ^41,47^. We have reported detecting short millisecond voltage pulses indicating a detection limit as low as 10 mV ^41^. The ability to optically measure voltage in a label-free manner opens new avenues in imaging bioelectrical signals at subcellular levels.

**Figure 1.**
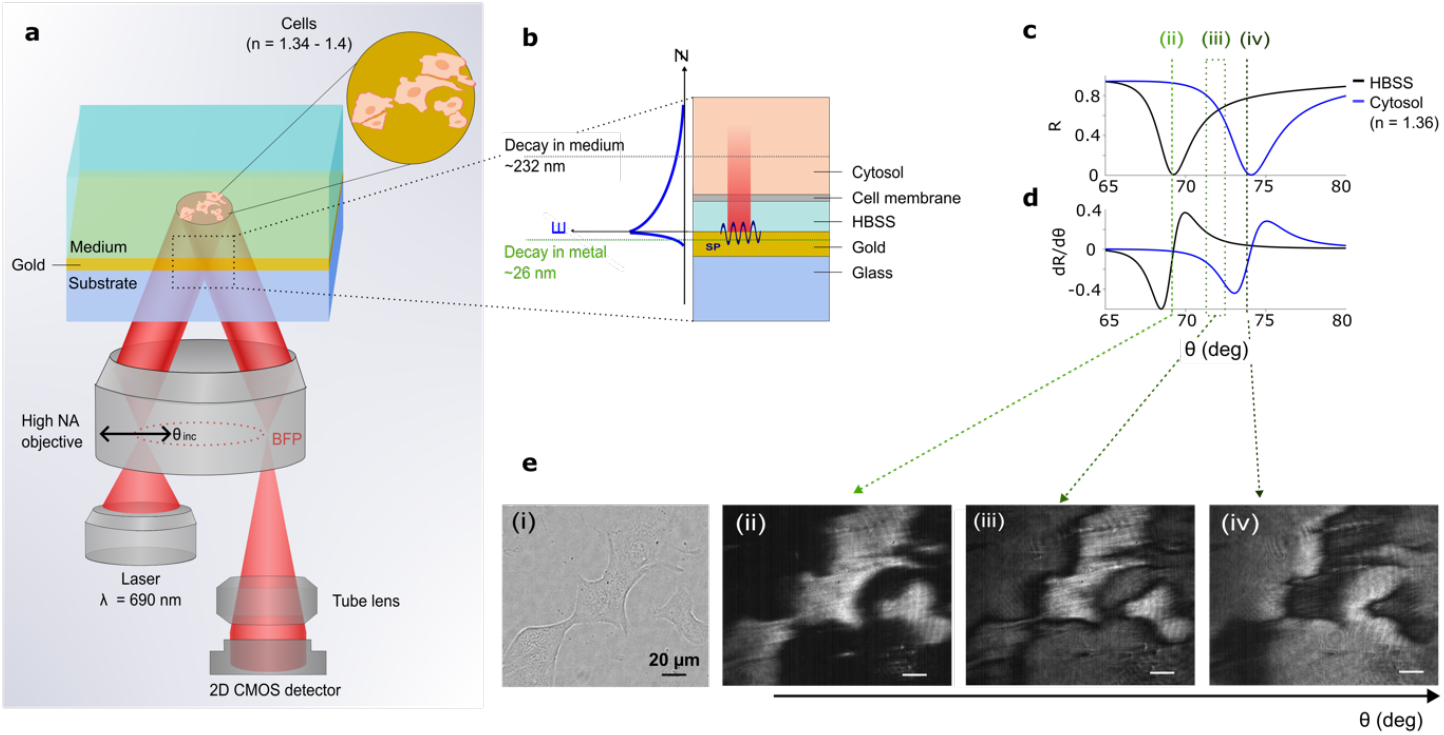
Surface Plasmon Resonance Microscopy (SPRM) of pancreatic beta-cells. **a** Schematic presenting the experimental setup for SPRM, depicting the layered structure of gold thin film on glass substrate with cells adhered to the gold surface in Hanks’ balanced salt solution (HBSS). A fiber-coupled laser (690 nm) is collimated before focusing on the back focal plane (BFP) of a high numerical aperture oil immersion objective to produce a collimated beam at the sample. The angle of illumination is varied by laterally scanning the focus on the BFP. The sample is imaged using a 2D CMOS pixelated detector. **b** A magnified view of the SPR sensor and cell interface showing the interface layers (glass, Au thin film of 50 nm, medium (HBSS), cell membrane of *c*.7 nm thickness, and cytosol), with an illustration of the penetration depth of SPs in both metal and dielectric media. This indicates sensitivity to the cell membrane and the proximal intra- and extracellular spaces. **c** SPR curves presenting the reflection coefficient for various angles of incidence, simulated for bare gold with HBSS and for the gold-cell interface respectively. **d** The corresponding first derivative of reflectivity with respect to the angle of incidence, showing the variations in the sensitivity of the measurement for optimising the angle of illumination. **e(i)**. Brightfield microscopy image of live MIN6 beta-cells cultured on PLL-modified Au thin film. **e**(**ii), e**(**iii)**, and **e**(**iv)** are the corresponding SPRM images at different angles of illumination. The angle of incidence is selected in region III, although this gives reduced sensitivity, it allows simultaneous tracking of cells and the extracellular regions where cells are not present on the sensor.

To demonstrate the SPRM applications in studying cell networks, the MIN6 immortalized cell line was chosen as an *in vitro* model of beta-cells and transferred to Hanks balanced salt solution (HBSS), as detailed in the Methods section, using poly-L-lysine (PLL) treated gold thin films as a substrate. To probe the cell sensor interface, we utilized the SPRM configuration illustrated in brief in Fig.1a and detailed in the Methods section. In this paper, we employed a widefield setup using a high numerical aperture oil immersion objective lens to excite SPs at an angle of incidence greater than that of the total internal reflection. The angle of incidence can be tuned by laterally translating a focus on the back focal plane of the objective lens.

To provide functional imaging capabilities, this configuration used an off-resonance angle of incidence for illumination where a subtle shift in resonance angle leads to a widefield change in the intensity of reflected light. Since, the aim of this study is to track both cells and their surrounding extracellular background, a careful selection of the angle of illumination will maximize the information retrieval. Therefore, we employed a transfer matrix-based simulation of the cell sensor interface to inform the choice of angle of incidence. Simulating the cell sensor interface, demonstrated in Fig.1b, produces the SPR curves corresponding to the cell and the background as presented in Fig.1c. A visual inspection shows that the points of high structural contrast (i.e., ii, and iv in Fig.1e) have the lowest functional sensitivities for both cells and background. Functional sensitivity is defined, here, as the ability to resolve small changes in the resonance angle or intensity due to dynamic living processes, which is realized by selecting an angle of illumination at a non-zero gradient of the SPR curve. The selection of an angle of incidence around the intersection of the two SPR curves allows signal dynamics to be probed with coverage of both cells and the surroundings. Nevertheless, this operating point (region III in Fig. 1c) does not necessarily provide the maximum sensitivities for both the cell and the background individually or the highest structural contrast. The cell-sensor interface is captured at different angles of incidence to illustrate the concept of functional sensitivity, as presented in Fig.1e.

We have previously presented a method based on mapping intensity gradient that can translate the change in intensity to the corresponding resonance angle shift ^40^. However, since for this study we are only interested in information propagation within the cell network, where connectivity is revealed using only correlations of phase and amplitudes, therefore, oscillation intensity was not quantitatively linked to the resonance angle shift. The next section shows the utilization of the SPRM approach to track the bioelectrical behaviour of pancreatic beta-cells.

### Ultra-high-density SPRM mapping of synchronized electrical oscillations

We report synchronized sub-second intensity oscillations in both cells and their extracellular background in the presence of HBSS supplemented with 10 mM glucose. We conjecture that the observed oscillations are linked to ionic currents associated with glucose-induced electrical signaling in pancreatic beta-cells, an idea which is validated in the next section. Coordinated electrical oscillations in pancreatic cells are well known to underpin the secretion of insulin by pancreatic islets ^44,48^. Figure 2 shows a cluster of three MIN6 cells, with bright-field and SPRM images in Figs. 2a and 2b, respectively.

**Figure 2.**
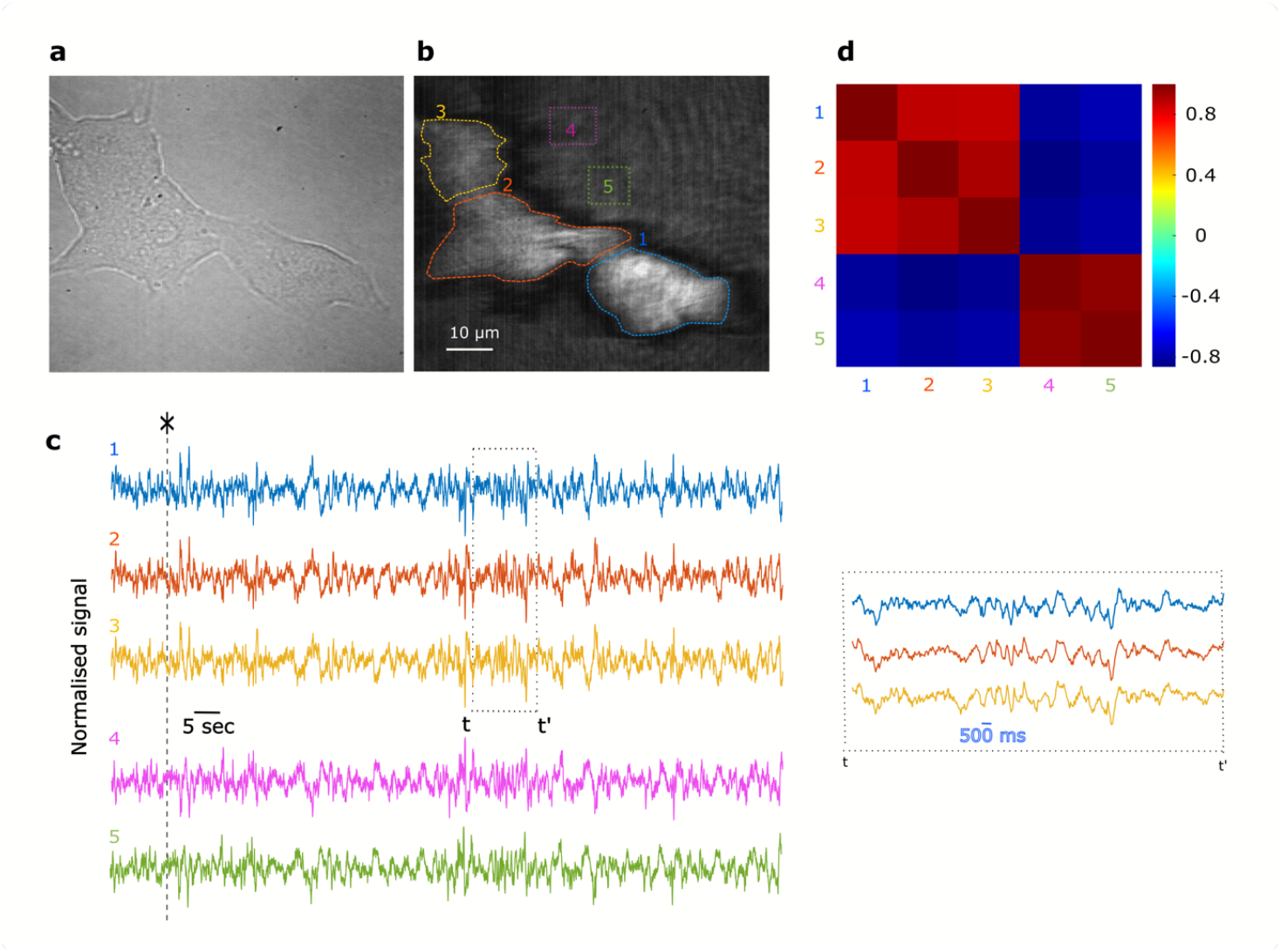
SPRM reveals correlated oscillations in pancreatic beta-cells. **a** Brightfield image of MIN6 cells cultured on PLL-modified Au thin film. **b** Corresponding SPRM image with five regions of interest highlighting cells (1, 2 and 3) and the extracellular background (4 and 5). **c** Time-resolved reflectivity recorded over 130 seconds in HBSS with 10 mM glucose for the five regions shown in **b**, inset shows a magnified view of a selected time window, indicated by t to t’, for the three cells which shows synchronised intensity oscillations. Traces appear synchronised but the background ROIs are anticorrelated. **d** Heat map displaying the correlation between signals extracted from ROIs 1 – 5 investigating signaling at the cellular ROIs (1-3) and the background ROIs (4, 5), where cells are not present.

Synchronous intensity oscillations are observed from regions of interest (ROIs) 1-5 where each ROI represents either a cell or the background. ROIs 1–3 (cells) display oscillations that appear anticorrelated with those from ROIs 4 and 5 (extracellular background), as shown in Fig. 2c. However, this anticorrelation is not physiological but results from the experimental conditions. In particular, the sample is illuminated with a collimated beam at a fixed angle of incidence, and due to refractive index differences, cells and their surroundings have resonance positions. As a result, the chosen angle of illumination leads to opposite intensity gradients between the cells and background (Figs. 1c and 1d), as discussed in the previous section. Therefore, correlated resonance position dynamics, between the cells and their background, lead to anticorrelated intensity changes.

This observation suggests that the SPR angle dynamics for both cells and the extracellular background are driven by the same biophysical process, which would lead to a correlated resonance angle shift. Pearson’s cross-correlation between these channels (i.e. ROIs 1 - 5) was computed and displayed in Fig. 2d showing two averages of 0.9067 ± 0.0367 and −0.8503±0.0218. This correlation map confirms that the oscillations are highly correlated within cells and extracellular background channels; however, there is anticorrelation between the two groups due to the SPR sensor transfer function.

The above findings are important from an electrophysiology perspective; the ability to obtain signals from both cells and the extracellular background is not possible with fluorescent based calcium and voltage imaging methods since they are only restricted to labelled cells. This new capability allows a thorough investigation of electrical signal propagation in pancreatic beta-cells and the surrounding medium. Although monitoring both cells and background is possible with microelectrode arrays, this is achieved at a reduced spatial resolution, of several micrometers enforced by the electrode spacing, even with high-density arrays ^12^. By contrast, SPRM provides an ultra-high-density recording capability, not previously possible, as demonstrated in Fig. 3. Here we show that these oscillations can be extracted from subcellular regions as small as 1 μm^2^. The spatial resolution of SPRM is diffraction limited in the direction perpendicular to SPs propagation while it is reduced to approximately 3 μm along the propagation direction ^49^. Therefore, while the method can report the global response from a ROI that averages over a whole cell, signals from subcellular regions can also be uncovered. To demonstrate this capability, the field of view was segmented into multiple channels, labelled (j, i) in Fig. 3a, each integrating the intensity within a 1 μm^2^ grid element. The small field of few of 85 μm x 91 μm is covered by more than 7700 channels, offering an exceptionally high spatial resolution. The correlation between the channels is clearly visible in Fig. 3b(i). Similarly, an anticorrelation between the cells and extracellular recordings is observed when comparing channels 4000 to 6000, where most of the MIN6 cells are located, to the remaining extracellular channels.

**Figure 3.**
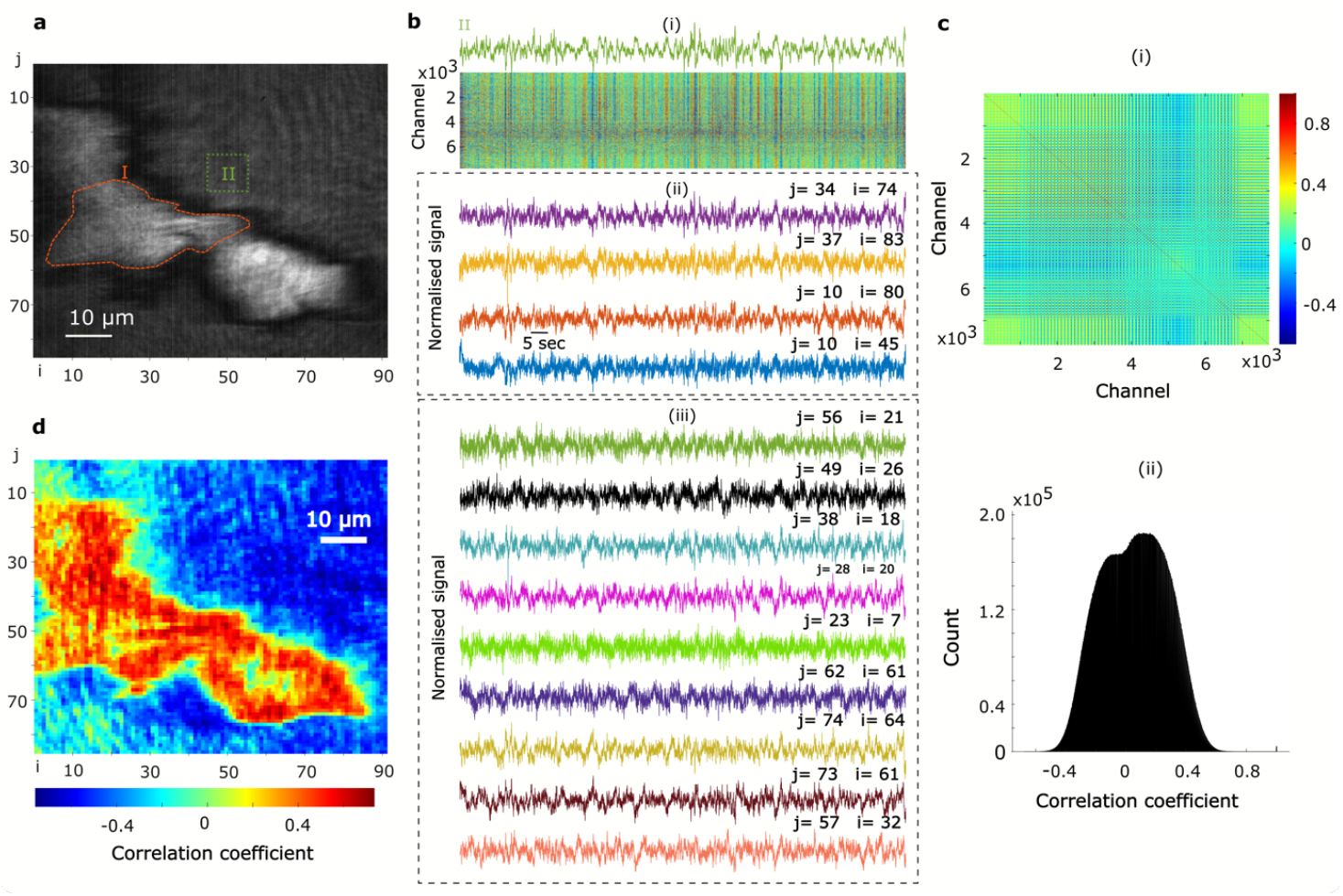
SPRM uncovers high-resolution sub-cellular oscillations in pancreatic beta-cells. **a** SPRM image highlighting two ROIs: I and II corresponding to a cell and the extracellular background, respectively. The image is segmented into multiple channels (pixels) of 1 μm^2^ labelled (i,j). **b** A stack of channels demonstrating ultra-high-density recording of electrical oscillations with a resolution of 1 μm_2_, shown in b(i). Exemplar traces are presented in b(ii) and b(iii) for the background and cell channels, respectively. **c** Pearson’s cross-correlation between the ultra-high-density channels, mapped in c(i) with the corresponding histogram depicted in c(ii). **d** The map illustrates the result of Pearson’s cross-correlation between a signal extracted by averaging over an exemplar individual cell (i.e. ROI II) and the local high-density sub-cell level signals (i.e. channels), revealing the spatial and temporal heterogeneity of the obtained signals.

The ultra-high-density recording capability can be leveraged to investigate the correlation and synchronization between subcellular signals. To achieve this, Pearson’s cross-correlation was computed for the channels described above and the correlation statistics are presented using the correlation map and the histogram in Figs. 3c(i) and 2c(ii), respectively. To rule out the contribution of a reduced signal-to-noise ratio at the subcellular regions leading to low correlations, we extracted the envelope of the oscillations using Hilbert transform before smoothing (methods sections). A similar correlation distribution to Pearson’s cross-correlation is observed eliminating the link between low signal-to-noise ratio and reduced correlations.

Next, we investigated the spatial origin and propagation of the observed intensity oscillations. Figures 2 and 3 illustrate global correlation between the MIN6 cells compared to local correlation statistics, obtained at the subcellular levels. Global correlations are defined as those between ROIs covering entire individual cells, while local correlations refer to those between subcellular signals. While the signals extracted by integrating over individual MIN6 cells (ROIs: 1-3) are highly correlated, as shown in Fig. 2(d), the subcellular signals show a lower correlation that is distributed over a range of −0.5 to 0.7, as presented in Figs. 3c(i) and 3c(ii). To visualise the subcellular correlations spatially, we calculated cross correlations between a global signal, integrated over a ROI covering an entire single cell (e.g. region I in Fig. 3a), and the signals extracted from the 1 μm^2^ sized channels shown in Fig. 3b(i). The resulting global-local cross correlation map is presented in Fig. 3d. A strong correlation is seen in the extracellular space surrounding cells, likely resulting from constructive interference between these cell-originated signals. On the other hand, the cellular recordings exhibit a relatively heterogeneous correlations, with higher correlations around the edge of the cells, in comparison to the central regions, indicating the spatial origin of the obtained signals. Representative time-series from cells and their background extracellular channels are presented in Figs. 3b(ii) and 3b(iii), the channel index i, j refers to the columns and rows that indicate the pixel location in the map in Fig. 3a.

### Synchronised intensity oscillations are suppressed via calcium channel blockers

To investigate the origin of the intensity oscillations, MIN6 cells were studied under three conditions consecutively: 1) baseline HBSS in the absence of glucose; (2) HBSS supplemented with 10 mM glucose; and 3) HBSS supplemented with 10 mM glucose and 40 μM of the calcium channel blocker nifedipine, as shown in Fig. 4. To compare the oscillation amplitudes, time series data from each cell were normalized to their respective standard deviation, calculated across the three experimental conditions. Cells exhibited spontaneous oscillations under baseline conditions, which increased in amplitude upon glucose exposure. However, when treated with nifedipine, the amplitude of these oscillations diminished, as observed in Figs. 4c and 4d.

**Figure 4:**
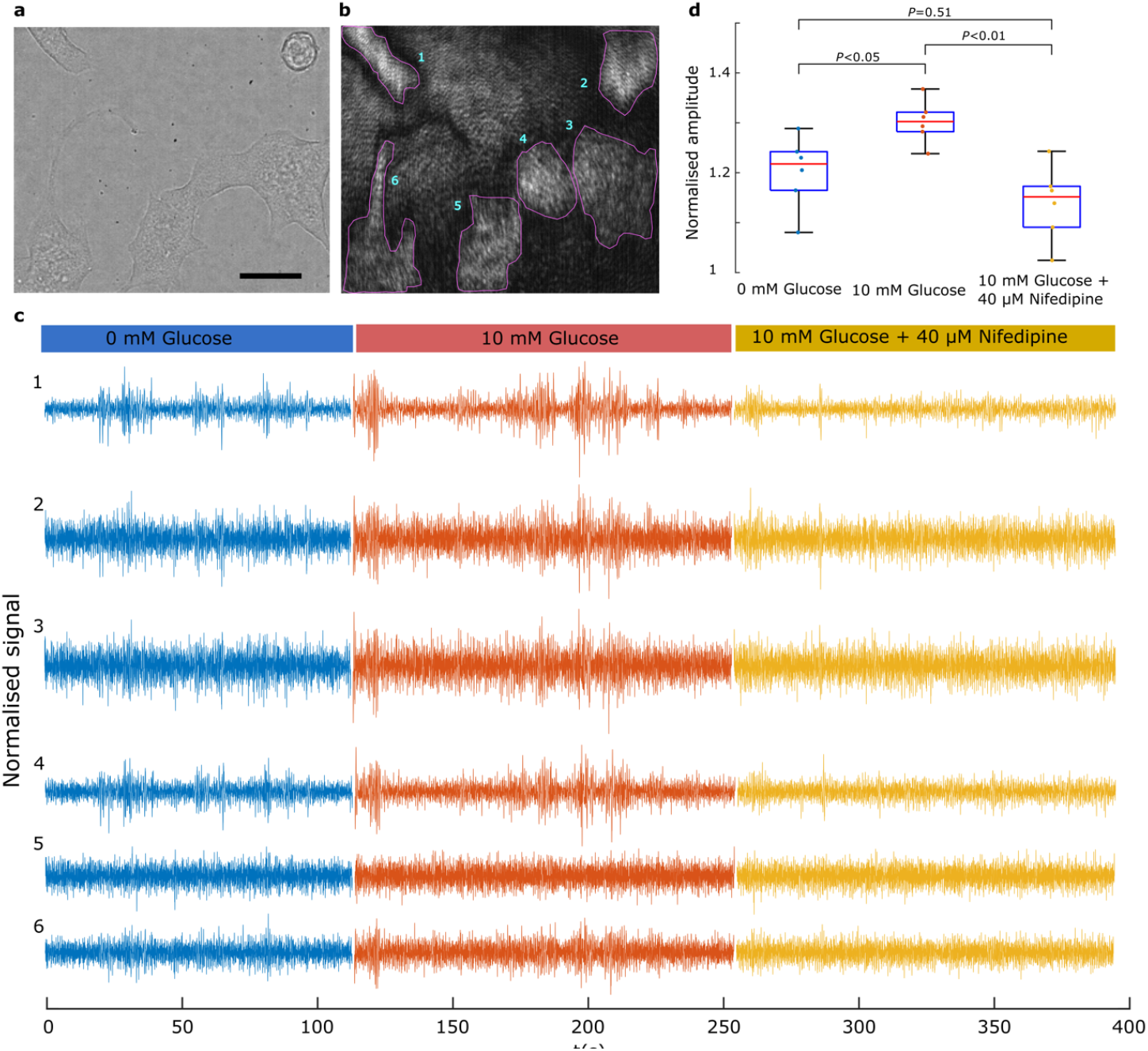
Synchronised network oscillations are suppressed in the presence of a calcium channel blocker. **a** Brightfield image of MIN6 cells cultured on PLL-modified Au thin film. **b** Corresponding SPRM image with 6 cellular regions of interest. **c** Time-series recordings from the six cellular ROIs presenting time-resolved reflectivity, under treatment with: 1) Hanks balanced salt solution (HBSS) without glucose; 2) HBSS supplemented with 10 mM glucose; and 3) HBSS supplemented with 10 mM glucose and 40 µM nifedipine. **d** Comparison of the effect of the three treatments on cells displaying the average amplitude profiles of the cells. Prior to identifying the amplitude profile, each signal was filtered between 0.1-15 Hz (see Methods) before standardization using the standard deviation over all the three recordings. (n=6; Bonferroni corrected *p* values).

The results, presented in Fig.4, indicate that the oscillations are linked to ionic dynamics across the cell membrane given the observed effect of the calcium channel blocker, nifedipine, which at the concentration used has well documented electrical activity blocking effects in pancreatic beta-cells ^50^. The spread of the signals beyond the regions of the cell electrode interface also supports the interpretation of the bioelectrical nature of the observed oscillations. One possible explanation of this latter phenomenon is that transmembrane ion dynamics modulate the charge at the double layer capacitor at the cell sensor interface leading to alteration of electron density in the metal, thus giving rise to the observed oscillations ^41^. Similarly, the perturbation of double-layer capacitor by transient increase or depletion of ion species could lead to changes in the refractive index in the dielectric layer adjacent to the metal. In the next section, we will demonstrate how these cells behave collectively as a network using techniques from computational graph theory to estimate connectivity across the network. Specifically, we use the phase-locking factor (PLF; explained in the next section) to estimate the influence of intensity oscillations in one cell to those in another.

### Functional connectivity in pancreatic cell networks

As discussed in the previous sections, dynamic SPRM of pancreatic beta-cells indicates an electrical origin for the synchronized oscillations observed among cells. Furthermore, the synchronization of these oscillations was quantitatively assessed using Pearson’s correlation coefficient. This observation is supported by the well-established knowledge of coordinated electrical signaling in pancreatic cell networks ^48,51^. This section presents a further investigation of connectivity between cells employing concepts from graph theory.

Functional connectivity was assessed using phase locking factor which was calculated for all combinations of the cell regions in the brightfield and the SPRM images in Fig. 5. Further details are provided in the methods section. The results for the three experimental conditions are presented using the connectivity matrices, depicted in Fig. 5c and respective directed network graphs in Fig. 5d. These adjacency matrices and the directed graphs are examples extracted from the first 100 seconds of each treatment. A feedforward connectivity is observed extending spatially from cell 1 towards cell 6, as indicated by the upper triangle in the adjacency matrices. Furthermore, the connectivity dynamics were investigated for each treatment, as presented in Fig. 5e. Both phase locking factor (PLF) and amplitude correlation coefficient (ACC) were calculated over a sliding 10-second window, stepped by one second. PLF was calculated as described in the methods section while ACC was computed by first applying the Hilbert transform to each time series followed by smoothing and then calculating undirected correlations between the resulting amplitudes. Examples of time series and their corresponding amplitude envelopes are presented in Figs. 5e(i) to 5e(iii).

**Figure 5:**
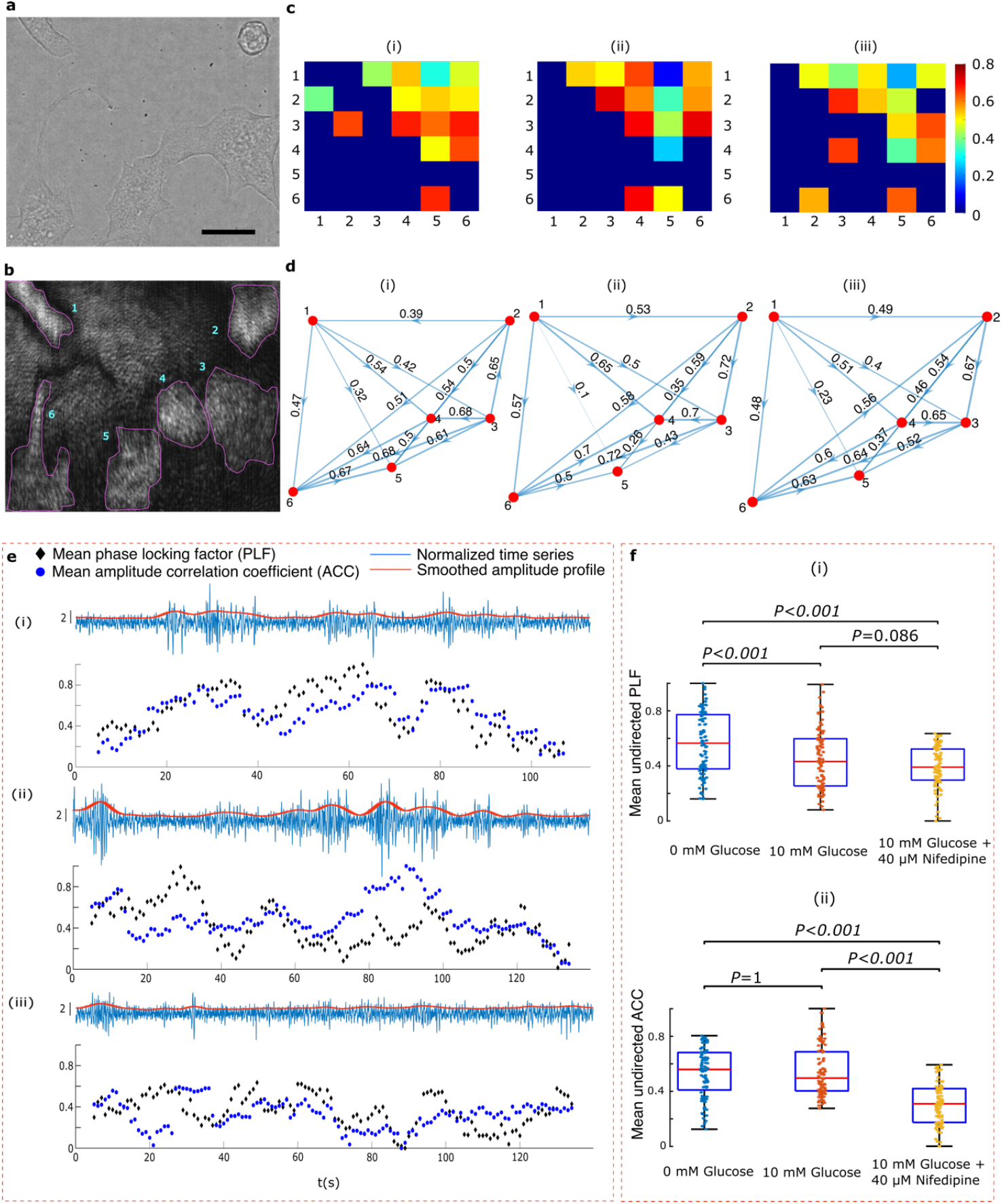
Network analysis. **a**, **b** Brightfield and SPRM images of MIN6 cells, respectively. **c** Connectivity matrices for the following conditions: baseline HBSS (**c.i**), HBSS supplemented with 10 mM glucose (**c.ii**) HBSS supplemented with 10 mM glucose and 40 μM nifedipine (**c.iii**). **d** Corresponding directed graphs represent cells ROIs as nodes, with edges (i.e. arrows) indicating patterns of directional connectivity and their associated weights. **e** Panels e(i) to e(iii) show examples of time series and their associated amplitude envelopes. Time-resolved connectivity, is presented for each of the above experimental conditions, measured via phase locking factor (PLF) and compared to amplitude correlation coefficient (ACC). PLF was calculated for a 10-second window with one second overlaps, for all cells and for each treatment. Similarly, ACC was computed by obtaining undirected correlation between the amplitude envelopes. **f** A boxplot showing the mean undirected PLF (i) and ACC (ii) extracted from the observations of their corresponding dynamics, for the six cells, presented in e(i) to e(iii). *P* values are Bonferroni corrected.

Figure 5f displays a boxplot of the observations of the mean PLF and the mean ACC reported in Fig. 5e. First, we observe, from Figs. 5f(i) and 5f(ii), a drop in both PLF and ACC upon exposure to nifedipine in comparison to baseline and glucose supplemented HBSS. Second, while ACC does not change before and after exposure to glucose, PLF drops significantly and upon exposure to glucose. This observation can be explained by inspecting Fig. 5e(ii). When cells are exposed to glucose, network dynamics show an anticorrelation between the ACC and PLF. While the amplitude envelopes are correlated during burst of activity, the cells undergo temporary, yet recoverable, phase decoherence (i.e. a drop in PLF), suggesting that connectivity drops temporarily during bursts. This observation indicates that cells burst in synchrony but, their individual spikes are incoherent ^52^. These results demonstrate the ability of the SPRM to reveal connectivity and the dynamics of a living cell network, in a label-free manner. The ability to observe and manipulate networks for extended durations is important to advance research in regenerative medicine.

## Discussion

The advancements in electronics, materials, nanotechnology and optics have led to bioelectric discoveries at multiple scales from single ion channels to macro brain circuits. However, method development research is still in demand to overcome the current technological limitations enabling label-free tracking of cell electrical signaling with high resolution and high spatial sampling. This work introduces a novel surface plasmonic method to study living cell networks in a minimally invasive manner, for the first time. Surface plasmonic imaging can reveal subtle changes in optical properties at the cell sensor interface, under the perturbation of ionic flow. Therefore, plasmonic imaging has created avenues for monitoring electrical properties of cells and biomolecules as well as electrical signaling in neurons ^16^ and cardiac myocytes ^17^. The results reported here demonstrate the ability to track electrical signaling from the subcellular level with a resolution of 1µm (Fig. 3) to a network level (Figs. 4 and 5), allowing a thorough investigation of information propagation within the network (Fig. 5). A small network of pancreatic beta-cells has been investigated to demonstrate the concept. The technique reveals highly correlated bioelectric signaling among cells, which is suppressed upon exposure to calcium channel blocker nifedipine (Fig. 4). When combined with graph theory, network structure and associated dynamics are uncovered. SPRM offers an ultra-high density recording capability (Fig. 3b(i)) and fine-grained imaging with a high spatial resolution down to 1 µm^2^ (Fig. 3d) that is not currently possible with the widely used microelectrode arrays. The optical readout offered by SPRM further eliminates the need for complex electrical wiring of recording channels. Furthermore, SPRM creates the opportunity to integrate complementary microscopy techniques to maximize the retrieval of bioelectric information and its morphological and biochemical correlates. Additionally, SPRM offers a high temporal resolution of several kHz, that is limited only by the speed of the 2D detectors and therefore can track fast voltage membrane dynamics. Several research directions are focussed at enhancing SPRM sensitivity and information retrieval through active plasmonics^53,54^, merging AI and plasmonics ^55^, and metamaterials ^56^, promising further advancement in SPRM imaging of electrical signaling.

Bioelectric signaling plays a key role in regulating living processes with cells forming complex bioelectric networks. For instance, functional neuronal networks are linked to behaviour, memory, cognition and bidirectional communications with internal and external environments. Decades of electrophysiological discoveries present several examples of bioelectric control of living processes such as homeostasis of blood glucose concentration. Corradiated electrical signaling in pancreatic beta-cells, within the islet of Langerhans, underlies the secretion of insulin sufficient to restore equilibrium. Pancreatic islets function as intelligent organoids, with integrated glucose sensing and insulin secretion (i.e., actuation) mechanisms to main homeostatic balance where organoid-level computations are orchestrated by electric signaling. This work leverages the SPRM recordings to investigate pancreatic beta-cell networks (Fig. 5). Phase locking factor, when applied to the obtained signals, shows a strong functional connectivity which is altered under exposure to the calcium channel blocker nifedipine. The technique can also resolve dynamics of networks connectivity and its relations to the amplitude (i.e. envelope) of oscillations.

Label-free monitoring of bioelectric networks promises long-term tracking of connectivity promising to expose the regeneration ^57^ and cognitive ^58^ correlates. Discoveries of leading research groups have shown the important role of bioelectricity in cancer metastasis ^59 8^, with cancer cells exhibiting fluctuation in membrane potential, similar to excitable cells, demonstrated with electrochromic voltage-sensitive dye ^7^. This study by Quicke *et al*, reported an intercellular correlation of V_m_ hyperpolarisation among cells, which hints to the importance of coordinated electrical communication. Furthermore, intercellular communication via depolarising potassium ion flow is linked to metabolic activity in bacterial biofilms ^9^. The ability to monitor living cell networks under different chemical, optical and bioelectronic stimuli creates a myriad of opportunities for engineering living networks with applications in drug discovery, bioelectronics^60,61^, regenerative medicine ^57^ and computing ^62,63^.

SPRM can track signal propagation beyond cells, as presented in Figs. 2 and 3. In this study, highly correlated oscillations beyond cells were observed that are likely produced due to constructive interference of cell-generated signals. This capability is important when studying long-range and extracellular communication between cells ^64^ and elucidating the effects of electric field on cell excitability ^65^, migration ^66^ and regeneration ^67^. The ability to image bioelectrical fields could enable a thorough investigation of different electrical pathways such as gap junction and extracellular routes in intercellular communications.

## Methods

### Simulations of SPR of cell-sensor interface

Transfer matrix method, a widely used method for studying light propagation in layered structures, was used to simulate the excitation of SPs given the variations in refractive index of the adhered sample. The cell sensor interface was studied using 1D model where light propagates through the following layers in the following order: glass (semi-infinite medium, n**=**1.5133), thin film of gold (50nm, n=0.13322+i3.9722)^68^, a cell medium gap between the cell membrane and the gold (thickness=50 nm, n=1.3350) ^69^, the cell membrane (thickness of 7.5 nm ^70^, n=1.4985^71^) and cytosol being a semi-infinite (n=1.36)^72^. The extracellular background with no cells present was modelled as glass, gold and semi-infinite cell medium having the same parameters mentioned above. SPR curve for both cells and their background were then calculated, at a wavelength of 690nm, by varying the angle of incidence in the range of (65° to 80° in steps of 0.01°) and calculating the resulting reflection coefficients.

### Cell culture

The mouse pancreatic cell line, MIN6 (Beta-TC-6, ATCC; CRL-11506), was maintained in high Glucose DMEM (Merck, D5671) supplemented with 10% FBS (Merck, F9665), 10 mM HEPES (Merck, R0887), 50 mg/ml penicillin and streptomycin (Merck, P0781) and 50 mM b-mercaptoethanol (Merck, M3148). Cells of passage numbers 35-42 were incubated in a humidified atmosphere containing 5% CO2 at 37°C. 24h prior to electrical measurement, MIN6 cells were transferred to RPMI 1640 media (Merck, R0883) supplemented with 11 mM glucose (Merck, G8644), 10% FBS, 10 mM HEPES, 50 mg/ml penicillin and streptomycin. Gold-coated or glass coverslips (diameter 22mm) were coated with 0.01% PLL (Merck, P04707) as described by the manufacturer to aid cell adhesion. Cells we seeded at a density of 10^5^ cells per coverslip in RPMI medium.

### SPR microscopy of pancreatic beta-cells

The label-free imaging of pancreatic beta-cell networks in this study was conducted using a Kretchmann-Raether configuration that is detailed in our previous study ^40^. Briefly, this setup is built around a high numerical aperture oil immersion objective lens (Nikon 60x NA 1.49). This objective lens is configured to illuminate the sample at a fixed angle of incidence using a collimated linearly polarized coherent light source (690nm, fiber-coupled). The fixed angle illumination is achieved by focusing light onto the back focal plane (BFP) of the objective lens. A tube lens is positioned at distance equal to the sum of its focal length and that of the objective lens, transforming the intensity of the BFP into an image of the sample, which is captured by a 2D camera (SV643M, EPIX, Inc., IL, US). The angle of incidence can be scanned by laterally translating the focus on BFP. For this purpose, the BFP of the objective is imaged to confirm the excitation of the SPs and optimize the measurement conditions.

After 24h growth, MIN6 cells seeded on gold-coated glass coverslips were removed from RPMI growth medium and placed in a static bath to obtain baseline recordings containing HBSS with the following composition (in mM): 137 NaCl, 5.6 KCl, 1.2 MgCl_2_, 2.6 CaCl_2_, 1.2 NaH_2_PO_4_, 4.2 NaHCO_3_ and 10 HEPES (pH 7.4 with NaOH). The bath was the replaced sequentially with HBSS supplemented with 10mM glucose prepared using a 1 M stock solution in water (10 µl/ml) followed by HBSS containing 40μM Nifedipine (Sigma), prepared from a 20 mM stock of Nifedipine in DiMethylSulfOxide. All experiments were conducted at 37°C.

Cells, under different treatments conditions, were monitored at frame rate of 100 Hz and the acquired videos were post-processed using Matlab. Time series were obtained by integrating the intensity over a manually defined region of interest marking a cell, which were used to investigate connectivity. Time series were also extracted from ROIs as small as 1μm. The signals are filtered using a bandpass zero-phase Butterworth filter described below. Correlation between the time series was calculated using Pearson’s correlation coefficient described in the next section.

### Network analysis

Network analysis was performed in MATLAB 2023B (MathWorks). Code is available to download from the GitHub repository: github.com/dgalvis/sprm_bcell_networks. Time series data were collected as averages of the SPRM signal over 6 manually identified cellular ROIs. These time series were filtered between 1-15 Hz using a 4^th^ order, zero-phase Butterworth filter. Unless otherwise noted, time series were then segmented into overlapping sliding windows of 10s – with a 1s increments (i.e., adjacent windows have a 9s overlap).

Phase locking factors (PLF) and amplitude-profile Pearson’s correlation coefficients (ACC) ^73,74^ were calculated for each of the 10s segments to study changes in pairwise phase and amplitude correlation (respectively) over time and conditions (i.e., 0 mM glucose, 10 mM glucose and 10 mM glucose supplemented with 40 μM nifedipine). Calculation of PLF and ACC require application of the Hilbert transform to produce a complex-valued signal

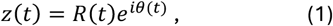

where *R*(*t*) is the amplitude profile and *θ*(*t*) is the phase profile for the signal.

The pairwise PLF between signals *i* and *j* is given by

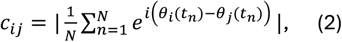

where *t*_*n*_ is the *n*th out of *N* sampling times. This method produces undirected networks, i.e., *c*_*ij*_ = *c*_*ji*_, and we consider the undirected networks unless otherwise noted. However, this method can be made into directed network method by retaining the values of *c*_*ij*_, if the angle of the complex number 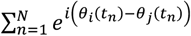 in eq2 is positive, otherwise, *c*_*ij*_ = 0.

The pairwise ACC between signals *i* and *j* is given by

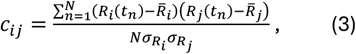

where 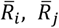 are the means and 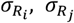 are the standard deviations of *R*_*i*_ and *R*_*j*_, respectively. In both methods, self-connections are excluded, i.e., *c*_*ii*_ = 0.

To mitigate against spurious connections due to finite-length time series, 99 surrogate datasets for each time window were generated using the iterative amplitude-adjusted Fourier transform (IAAFT) method ^75,76^. This method produces surrogate datasets that preserve autocorrelation whilst removing pairwise cross-correlations in the original signals. Using this method, a connection *c*_*ij*_ is rejected if it does not exceed the 95% level of significance (i.e., if *c*_*ij*_ > *s*_*ij*_ in less than 95% of cases, where *s*_*ij*_ is the corresponding connection in a surrogate dataset). Where a connection is rejected, the connection is set to zero, i.e., *c*_*ij*_ = 0. Figure 5e shows the average connection strength over the network ⟨c_ij⟩ for each time window. Average ACC and PLF values were renormalized over all time windows and conditions such that the minimal and maximal values were 0 and 1, respectively.

## Acknowledgments

This work was supported by the Engineering and Physical Sciences Research Council [EP/M50810X/1, EP/X018024/1], UKRI [MR/X034240/1], and the University of Nottingham. SA acknowledges the financial support of Nottingham Research Fellowship. DG acknowledges financial support from the University of Birmingham Dynamic Investment Fund.

